# Signaling specificity and kinetics of the human metabotropic glutamate receptors

**DOI:** 10.1101/2023.07.24.550373

**Authors:** Tyler W. McCullock, Loren P. Cardani, Paul J. Kammermeier

**Author notes:** All authors have approved submission of this manuscript. This work has not been accepted or published elsewhere. Authors declare no competing interests.

## Abstract

Metabotropic glutamate receptors (mGluRs) are obligate dimer G protein coupled receptors that can all function as homodimers. Here, each mGluR homodimer was examined for its G protein coupling profile using a BRET based assay that detects the interaction between a split YFP-tagged Gβ_1_*γ*_2_ and a Nanoluc tagged free Gβ*γ* sensor, MAS-GRK3-ct-NLuc with 14 specific G⍺ proteins heterologously expressed, representing each family. Canonically, the group II and III mGluRs (2&3, and 4, 6, 7&8, respectively) are thought to couple to G_i/o_ exclusively. In addition, the group I mGluRs (1&5) are known to couple to the G_q/11_ family, and generally thought to also couple to the PTX-sensitive G_i/o_ family; some reports have suggested G_s_ coupling is possible as cAMP elevations have been noted. In this study, coupling was observed with all 8 mGluRs through the G_i/o_ proteins, and only mGluR1&5 through G_q/11_, and perhaps surprisingly, not G_14_. None activated any G_s_ protein. Interestingly, coupling was seen with the group I and II, but not the group III mGluRs to G_16_. Slow but significant coupling to G_z_ was also seen with the group II receptors.

## Introduction

Metabotropic glutamate receptors (mGluRs) are class C G protein coupled receptors, and consist of 8 members (mGluR1-8), organized by sequence homology, signaling effectors, and general localization (*1*) . The Group I mGluRs include mGluR1 and mGluR5, are dual-coupled through Gα_q_ and Gα_i/o_ (*2–5*), and exhibit post-synaptic expression (*6*) in the nervous system, and there have been reports of cAMP accumulation in response to mGluR1 and mGluR5 activation giving rise to speculation of G_s_ coupling by these receptors (*7–10*). The Group II mGluRs consist of mGluR2 and mGluR3 and are thought to couple exclusively to the Gα_i/o_ pathway (*11*). These receptors can be found at either the pre- or post-synapse participating in cAMP based synaptic plasticity as well as acting as auto receptors via Gβ*γ* to limit the amount of glutamate released during action potentials (*12*). The Group III mGluRs consist of mGluR4, mGluR6, mGluR7, and mGluR8. These receptors also believed to solely couple to G_i/o_ signaling pathways (*11*). mGluRs 4, 7, and 8 are typically found acting as auto receptors on the pre-synaptic terminus (*13*) while mGluR6 expresses exclusively post-synaptically in retinal ON bipolar cells (*14*). Interestingly though, mGluR7 is only poorly responsive to millimolar concentrations of glutamate and no other native agonist has been identified for it (*15–17*). While the G protein coupling tendencies of the mGluRs is generally known, a comprehensive assessment of mGluR-G protein coupling has not been published, although some studies have examined the coupling of representative members of each group (*18*).

Due to their widespread expression in the nervous system, mGluRs participate in many neuronal physiological processes and pathophysiological behaviors. For this reason, mGluRs have been considered potential therapeutic targets for a wide range of pathologies including addiction, epilepsy, schizophrenia, and Parkinson’s disease (*19*). Here we utilize optimized adaptations of state of the art bioluminescence resonance energy transfer (BRET) assays to assess the G protein signaling of each member of the mGluR family as homodimers in detail in HEK293T cells.

Our results indicate that all members of the mGluR family can activate members of the G⍺_i/o_ family, while only the group I receptors, mGluR1 &5, couple to G_q_ and G_11_. Interestingly, we observe coupling through G_16_ through mGluRs 1, 5, 2, and 3 only, although coupling with mGluR3 was quite weak. None of the mGluRs exhibited coupling to the G_12/13_ or G_S_ families, even mGluR5 in the presence of the positive allosteric modulator VU0424465, which had been reported to promote G⍺_s_ coupling (*8*). In addition, the kinetics and potencies of each mGluR coupling to their corresponding G⍺ proteins were also examined. In general, the group II mGluRs appeared to be the most efficient activators of G⍺ proteins. The group II and III receptors activated the G_o_ proteins with the highest potency, while the Group I receptors most potently activated G_q_.

## Results

### Optimizing the NanoBRET assay for mGluR signaling

Our goal was to comprehensively examine receptor-G protein coupling profiles of each of the 8 human mGluR homodimers (Fig. 1A) with members of each family of G proteins. To accomplish this, we employed an optimized version of a Gβ*γ* based BRET assay (*20, 21*) that detects the interaction between the Gβ*γ* binding region of GRK3 fused to NanoLuc (NLuc) on its C-terminus and to a myristic acid sequence on its N-terminus (MAS-GRK3-NLuc), and a complemented YFP-tagged Gβ*γ* that is sequestered when inactive by a heterologously expressed G⍺ (“NanoBRET”; Fig1B). Each construct (see Materials and Methods), along with the indicated receptor was expressed in HEK293T cells (Fig. 1B). However, because HEK cells secrete micromolar concentrations of glutamate into the extracellular space (*22*), assay conditions needed to be optimized compared with those originally published (*21*), to reduce ambient glutamate levels that could potentially produce basal activation of mGluRs, which could produce high apparent basal BRET signals and reduce the observed ΔBRET, as shown in Fig. 1C, using mGluR2 and G_oA_, which shows high basal BRET signals and small ΔBRET upon application of 1 mM glutamate. The dramatic reduction in BRET signal in this experiment when the competitive antagonist LY341495 was applied with or without glutamate (Fig. 1C, *red* and *blue,* respectively) demonstrates that the high basal BRET signal was likely due to ambient glutamate in the well. To address the elevated glutamate in the bath, a combination of amino acid transporter expression (*23, 24*), washing, and timing of the experiments was used. Conditions were optimized to 1) reduce the basal BRET ratio, 2) reduce the responsiveness to a pan-mGluR antagonist (LY341495) in the absence of exogenous agonist, and 3) maximize the ΔBRET signal generated by glutamate application. Fig. 1D, shows responses of mGluR2/G_oA_ to the optimized protocol (also see Materials and Methods). Note the lower basal BRET value, the reduced effect of antagonist (*red*), and the strengthened ΔBRET signal upon application of 1 mM glutamate (*green*). A summary of the basal BRET levels (Fig. 1E), the change in BRET with LY341495 (Fig. 1F), and the ΔBRET upon glutamate application (Fig. 1G) or glutamate +LY34 (Fig. 1H) is also shown using the standard (red) and optimized (gray) NanoBRET protocols. Effects of null receptor conditions are also shown (“-/oA,” and “-/q”), illustrating that no responses are seen in the absence of heterologous mGluR expression. To be certain that each G⍺ protein expressed in the NanoBRET assays was expressed and functional, control experiments were performed with receptors that canonically couple to all of the G protein families to be tested. For these experiments, we used the H1 histamine receptor (hH1R; G_i/o_ and G_q/11_), the lysophosphatidic acid receptor 2A (mLPA2R; G_i/o_, G_q/11_, and G_12/13_) (*18*), and the D5 dopamine receptor (hD5R; the G_s_ family). The combined results of these experiments (Fig. S1) show that positive results can be obtained with each of the G⍺ proteins expressed in our NanoBRET assays.

**Figure 1.**
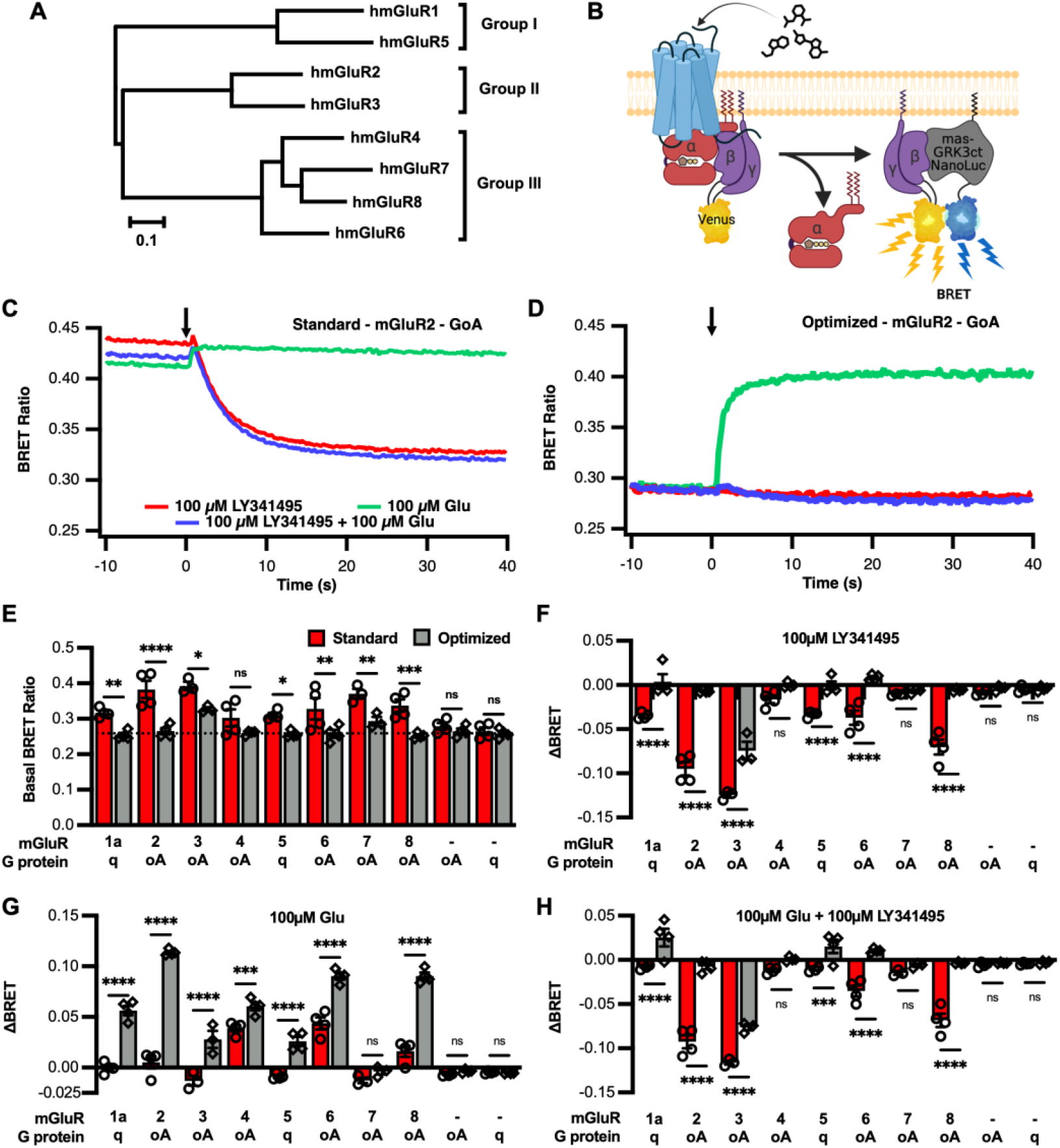
Evaluation of the mGluR optimized protocol for all human mGluRs. A, Dendrogram illustrating homology of all 8 human mGluRs and their distribution into three groups. B, Schematic illustrating the setup of the NanoBRET system. C-D, Example protocol of cells expressing mGluR2-G_oA_ prepared under the standard protocol (C) or mGluR optimized protocol (D) to 100 μM glutamate (green), 100 μM LY341495 (red), or the combination of the two (blue). The stimulus was delivered at time = 0 as indicated by the arrow. Summary data describing the basal BRET ratio (E), the LY341495 response (F), the glutamate response (G), and responses to Glu+LY34 (H) of all 8 mGluRs in the experiments as shown in C&D. Bars describe the average ± SEM. Statistics show the results of a two-way ANOVA with Holm-Šídák post hoc test, * = P<0.05, ** = P<0.005, *** = P<0.0005, and **** = P<0.0001.

### Group I mGluR profiles

To begin to assess mGluR-G protein coupling, each group I mGluR (1&5) was expressed in combination with the optimized NanoBRET system with a panel of 14 G⍺ proteins spanning all 4 major families. In each experiment, 1 mM glutamate was added at 0 seconds. Fig. 2A shows averaged, time resolved ΔBRET traces for 14 G⍺ proteins in cells expressing mGluR1, which responded to each member of the G_i/o_ family except for G_z_, and as expected responded to the G_q/11_ family of G⍺ proteins with the notable exception of G_14_. G_q_ appeared to be activated with the highest efficacy (Fig. 2A&B), while G_11_, G_oA_ and G_oB_ also responded with high efficacy and G_i1-3_, and G_16_ responded somewhat strongly as well. No responses were observed indicating coupling of mGluR1 with the G_s_ family members S_short_, S_long_, or Olf (Fig. 2A&B). Kinetics of activation of each responding G protein were also assessed by calculating the initial rate of activation for each (Fig. 2C; also see Materials and Methods).

**Figure 2.**
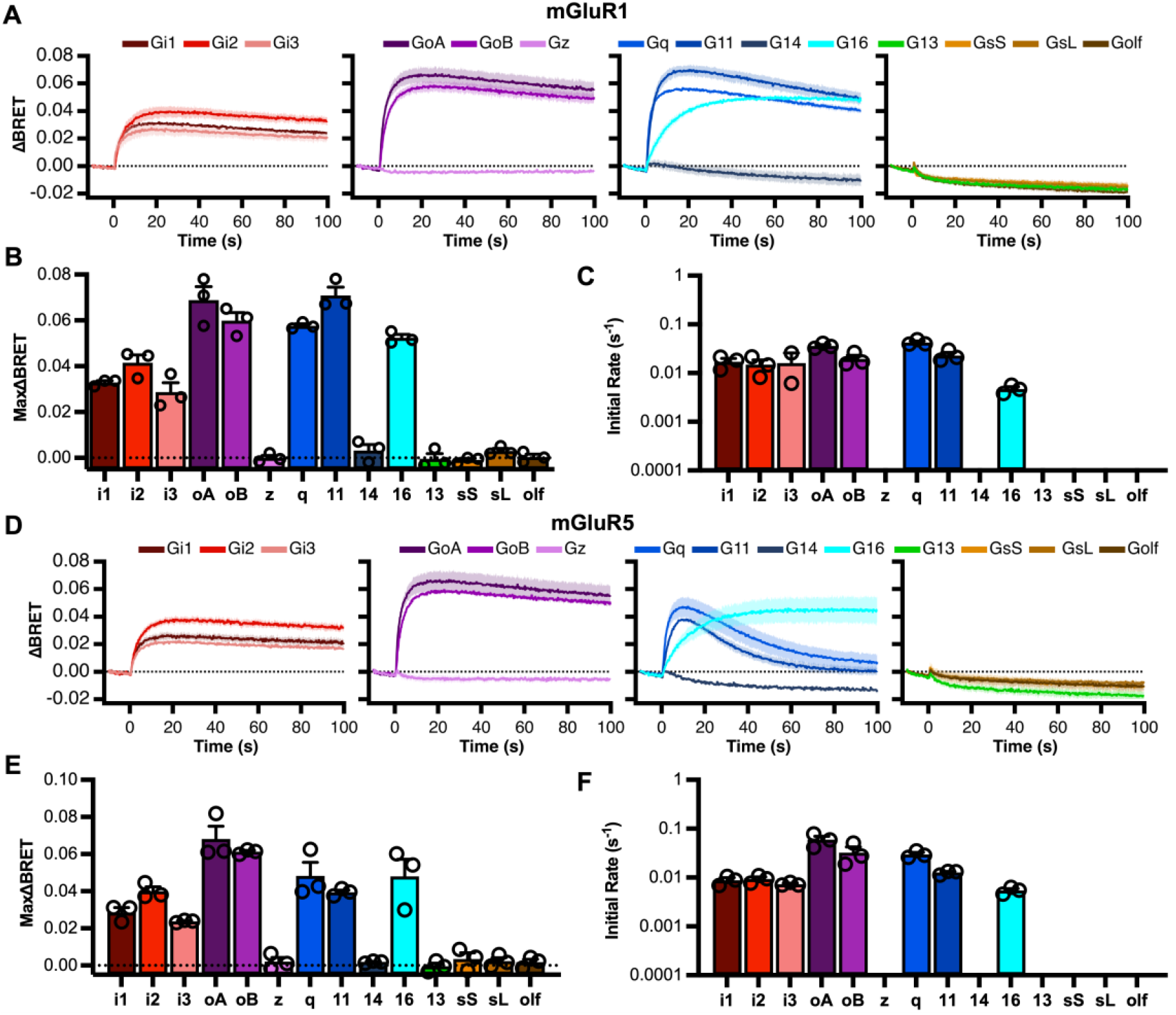
Group I mGluR signaling profiles. Signaling profiles of mGluR1 (A-C), and mGluR5 (D-F), through a panel of 14 G⍺ proteins in response to 1 mM glutamate. Each stimulus was delivered at time = 0. The dark solid line indicates the average response of 3 biologic replicates and the light shading indicates the SEM for each trace. Maximum ΔBRET induced by glutamate (B&E) and initial rates (C&F) for mGluR1 and 5, respectively are displayed as the average ± SEM. Raw measurements for each replicate are shown as open circles (○) in each bar graph.

The profile of mGluR5 was qualitatively similar to mGluR1 but with a notable apparent desensitization of responses to G_q_ and G_11_ (Fig. 2D), which has been documented previously (*25, 26*). This desensitization may have hindered measurement of the full efficacy of these responses. As such, mGluR5 coupled most strongly to G_oa_ (Fig. 2D-F).

Because previous reports have indicated that group I mGluRs can initiate cAMP accumulation and may therefore couple to G_s_ proteins (*7*), and a recent study has suggested that purified, truncated, mGluR5 is capable of activating G_s_ in the presence of the agonistic positive allosteric modulatior (PAM) VU0424465 (VU042) (*8*), we re-examined the coupling profile of mGluR5 in the presence of glutamate alone, VU042 alone, and glutamate + VU042 together (Fig. S2). While we observed some differences in maximum BRET with glutamate compared with glutamate + VU042 when coupling to G_i1_, G_i3_, and G_oB_ (Fig. S2), we did not observe mGluR5 coupling to any G_s_ family member in any condition. Together, these data confirm that the group I mGluRs are dual coupled receptors that can couple with high efficacy to the G_i/o_ and G_q/11_ families.

### Group II profiles

When assayed in the optimized NanoBRET system, mGluR2 yielded detectable coupling to each member of the G_i/o_ family (G_i1-3_, G_oa_, G_oB_, G_z_) as well as the promiscuous G_16_. The kinetics of G_z_ and G_16_ activation by mGluR2 were dramatically slower, however. Note that the S1 subunit of pertussis toxin (PTX) was co-expressed (*27*) with each PTX-insensitive G⍺ protein, including G_z_, to prevent a small but detectable signal presumably carried by endogenous G_i/o_ proteins in these and all subsequent experiments. Similar results were seen with the other group II member, mGluR3 (Fig. 3), although because of the high potency of this receptor, and its apparent consequent basal activation leading to high basal BRET ratios even under optimized conditions (Fig. 1F), it was necessary to reduce extracellular Cl^-^ levels to right-shift mGluR3 potency (*28*), to obtain meaningful data (see below). Re-assaying mGluR2’s signaling profile under low Cl^-^ conditions showed a similar coupling profile as in the standard High Cl^-^ buffer, which justifies using this method for mGluR3 measurements. Fig. S3 shows the maximum BRET amplitude and activation kinetics with mGluR2 (Fig. S3D) as well as correlations (Fig. S3E&F) of these measurements for each G protein in low and high Cl^-^ conditions.

**Figure 3.**
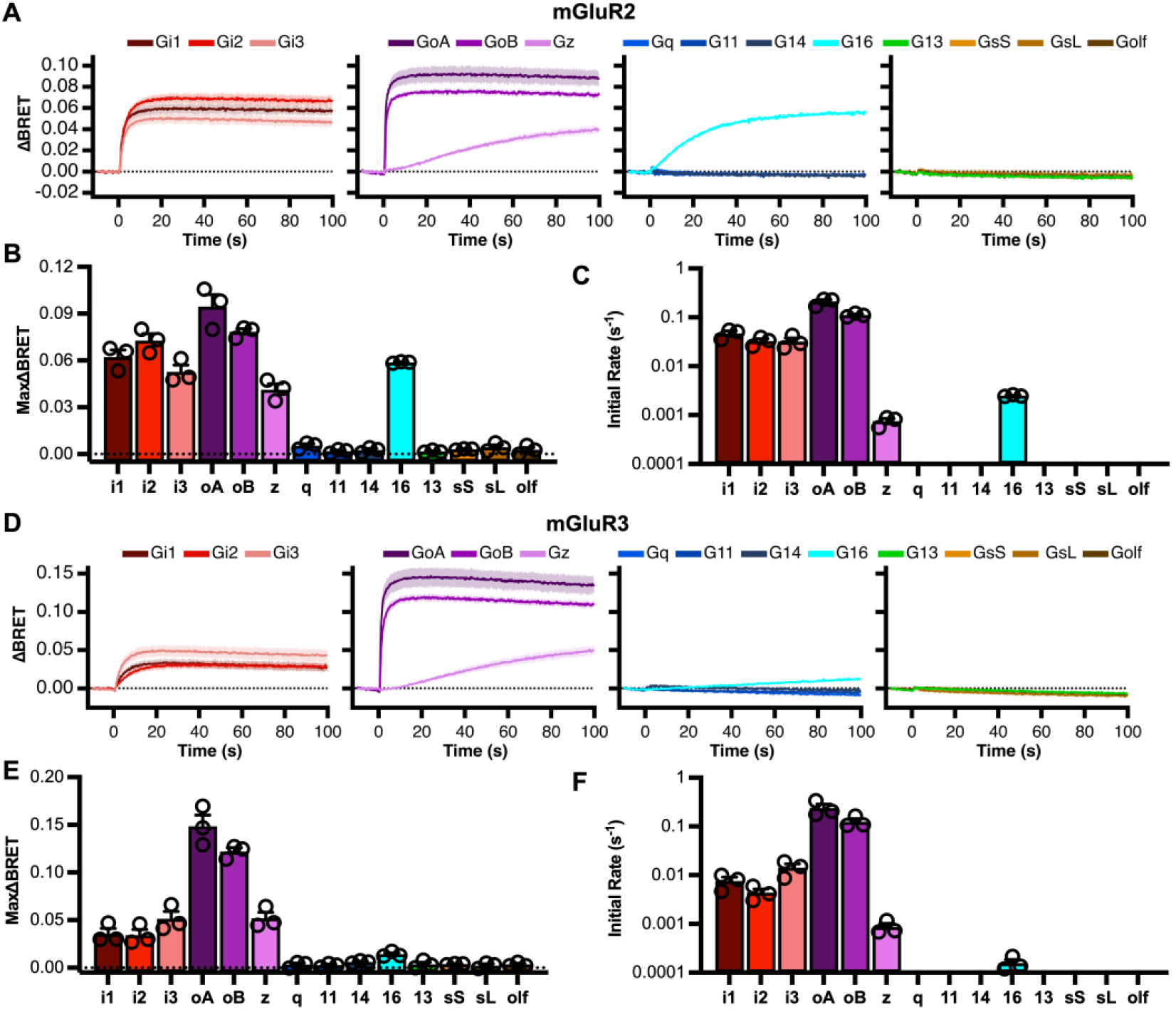
Group II mGluR signaling profiles. Group II mGluR signaling profiles. Signaling profiles of mGluR2 (A-C), and mGluR3 (D-F), through a panel of 14 G⍺ proteins in response to 1 mM glutamate. Each stimulus was delivered at time = 0. The dark solid line indicates the average response of 3 biologic replicates and the light shading indicates the SEM for each trace. Maximum ΔBRET induced by glutamate (B&E) and initial rates (C&F) for mGluR1 and 5, respectively are displayed as the average ± SEM. Raw measurements for each replicate are shown as open circles (○) in each bar graph.

Correlations show a consistent shift to higher potencies in low Cl^-^ but maintain the slopes, indicating that the rank order of G protein coupling remains unaltered. Thus, examining the full profile of mGluR3 in low Cl^-^ reveals that like mGluR2, mGluR3 couples almost exclusively to the G_i/o_ family, although favors coupling to G_oA_ and G_oB_ vs. G_i_ proteins (Fig. 3B). Like with mGluR2, mGluR3 showed a weaker, slower activation of G_z_, but in contrast, no detectable coupling to G_16_. As expected, neither group II receptor coupled to any members of the G_q_ or G_s_ families.

### Group III profiles

G protein coupling was also similarly assessed with the group III receptors mGluR4-8 (Figs. 4&5). As with mGluR2, each of the G_i/o_ family members were activated by mGluR4, although no detectable activation of G_16_ was observed, and coupling to G_Z_ was very weak (Fig. 4). Similar profiles were observed with all of the group III mGluRs, mGluR6 (Fig. 4), as well as 7 and 8 (Fig. 5), confirming that these receptors are exclusively coupled to the G_i/o_ family, and demonstrating variable G_Z_ coupling within the group III mGluRs. In this group, mGluR7 was somewhat anomalous, only showing weak coupling to G_oA_ and G_oB_ (Fig. 5A&B). Likely due to its high constitutive activity (*29*) and very low potency, this receptor exhibited a high basal BRET signal (Fig. 1), and required 10 mM glutamate to detect its comparatively poor activation (Fig. 5). It is possible mGluR7 may be capable of coupling to other G proteins, but our assay would only sense this if more efficient activation of mGluR7 could be achieved. In apparent contrast to the GABA_B_ receptor (*30*), none of the mGluRs showed detectable activation through G_13_, and none showed activation of G_14_ or the G_s_ family.

**Figure 4.**
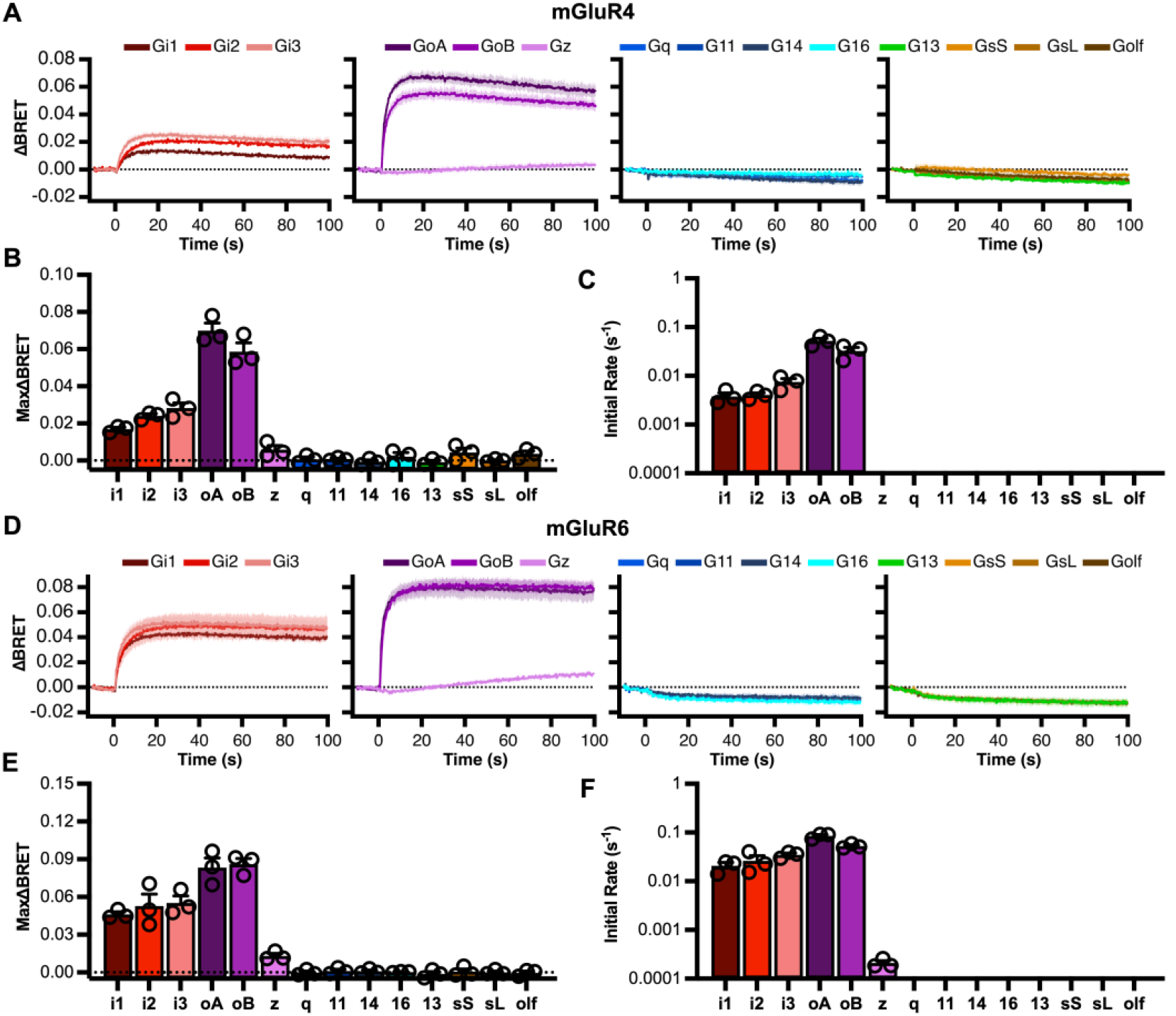
Group III (mGluR4&6) signaling profiles. Group III mGluR signaling profiles. Signaling profiles of mGluR4 (A-C), and mGluR6 (D-F), through a panel of 14 G⍺ proteins in response to 1 mM glutamate. Each stimulus was delivered at time = 0. The dark solid line indicates the average response of 3 biologic replicates and the light shading indicates the SEM for each trace. Maximum ΔBRET induced by glutamate (B&E) and initial rates (C&F) for mGluR1 and 5, respectively are displayed as the average ± SEM. Raw measurements for each replicate are shown as open circles (○) in each bar graph.

**Figure 5.**
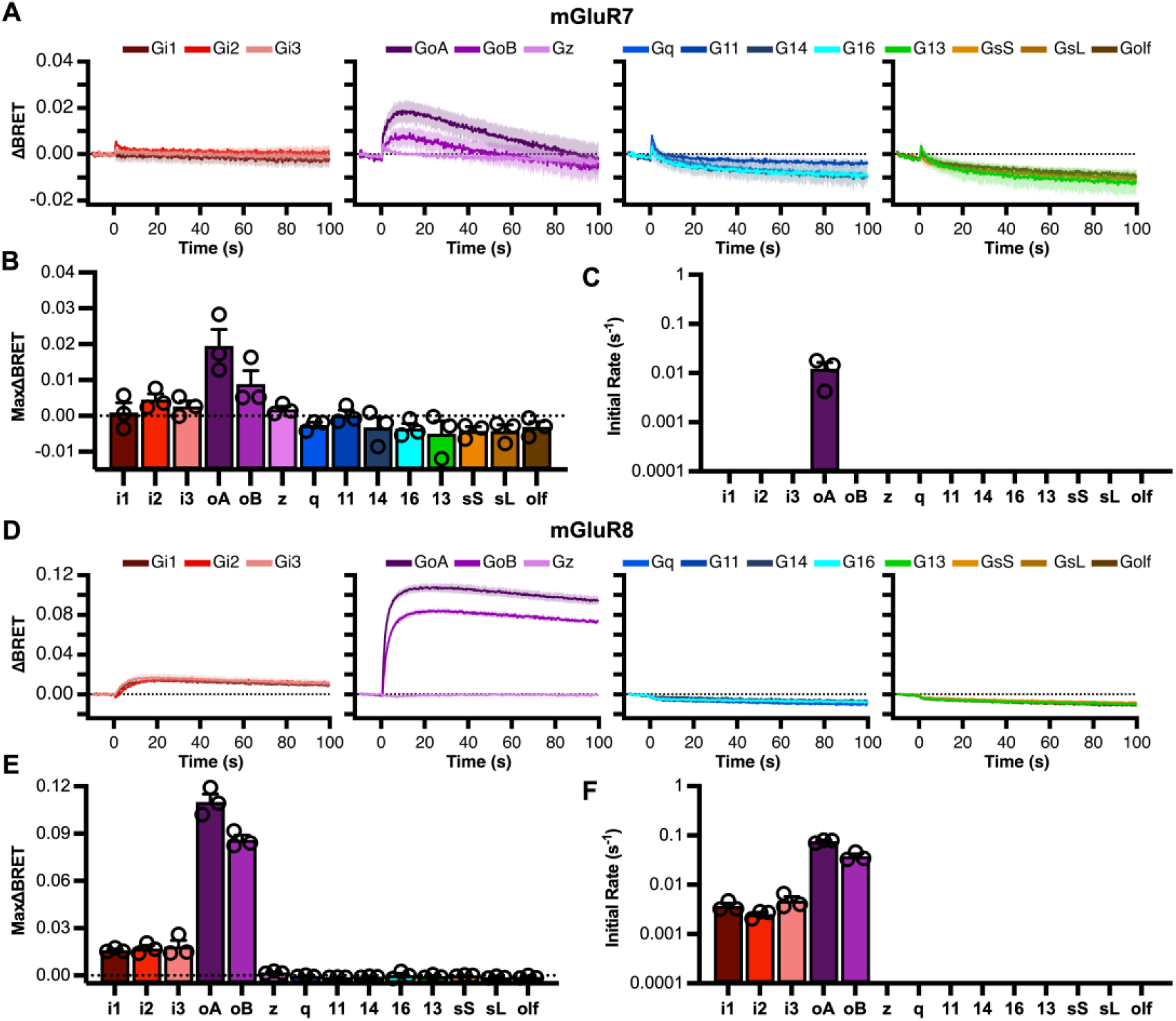
Group III (mGluR7&8) signaling profiles. Group III mGluR signaling profiles. Signaling profiles of mGluR7 (A-C), and mGluR8 (D-F), through a panel of 14 G⍺ proteins in response to 1 mM glutamate. Each stimulus was delivered at time = 0. The dark solid line indicates the average response of 3 biologic replicates and the light shading indicates the SEM for each trace. Maximum ΔBRET induced by glutamate (B&E) and initial rates (C&F) for mGluR1 and 5, respectively are displayed as the average ± SEM. Raw measurements for each replicate are shown as open circles (○) in each bar graph.

Because of the need to test mGluR3 responses in low Cl^-^ as described above, dose response curves were generated for each receptor (except mGluR7) in normal and low Cl^-^ (Fig. S4). Full dose responses were generated with the highest potency G⍺ protein with each receptor (G_q_ for the group I receptors and G_oA_ for group II and III) in high (144 mM) and low (7 mM) Cl^-^. Interestingly, reducing the Cl^-^ concentration resulted in a right shift in the potency of every receptor tested. Rescue of the mGluR3 responses in low Cl^-^ is consistent with the interpretation that the low levels of basal extracellular glutamate present in these experiments is enough to activate and desensitize these receptors (*31*) when measured at high Cl^-^, but low Cl^-^ shifts the potency such that ambient glutamate is below the threshold of activation, and therefore avoiding high basal activity and desensitization of this receptor in high Cl^-^. Net effects of Cl^-^ changes on potency (Fig. S4B) and efficacy (Fig. S4C) are also shown. Finally, Fig. S4D&E illustrate that the rank order of G protein potency with mGluR2 is unaffected by the change in [Cl^-^], suggesting that low Cl^-^ remains a reasonable modification to measure responses through mGluR3.

### Potency of mGluR homodimer signaling through different G⍺ proteins with glutamate

To assess the potency of each mGluR through each identified G⍺ protein signaling partner, we employed the NanoBRET system at a range of glutamate concentrations (Fig. 6). For each receptor, dose response data was only obtained with G⍺ proteins that showed significant responses in the profiling assay (Figs. 2-5). The group I mGluRs, mGluR1&5, were the only mGluRs that showed responses with G_q_ and G_11_, and both of these receptors responded with the highest potency with G_q_ activation, which was slightly higher than G_11_ in each case (Fig. 6A). In both cases, G_q_ and G_11_ signaling was also slightly more potent than signaling through G_i/o_ proteins, which all showed very similar EC_50_ values. The group II mGluRs showed a clear preference for G_oA_ and G_oB_ (Fig. 6A) and mGluR2 showed intermediate potency with G_i1-3_, and lowest potency activation of G_z_ and G_16_, while mGluR3 (in low Cl^-^) exhibited similar responses to G_i1-3_, z, and G_16_. In general, the group III receptors also activated G_oA_ and G_oB_ with the highest potency, followed by G_i1-3_, and finally G_z_ (mGluRs4&6; Fig. 6). Due to the very low potency for mGluR7 action through most G proteins, dose response curves could only be obtained with G_oA_, and these could not be tested to saturation due to solubility as well as osmolarity issues at very high glutamate concentrations. Heat maps summarizing the EC_50_ values for each receptor with each G⍺ protein tested are shown in Fig. 6B, and the Hill coefficients for each condition tested are shown in Fig. 6C. Note that efficacy for this data set was normalized for each pathway to allow for easier comparison of potency, but in all cases, efficacy was similar to the Max ΔBRET values shown in Figs 2-5.

**Figure 6.**
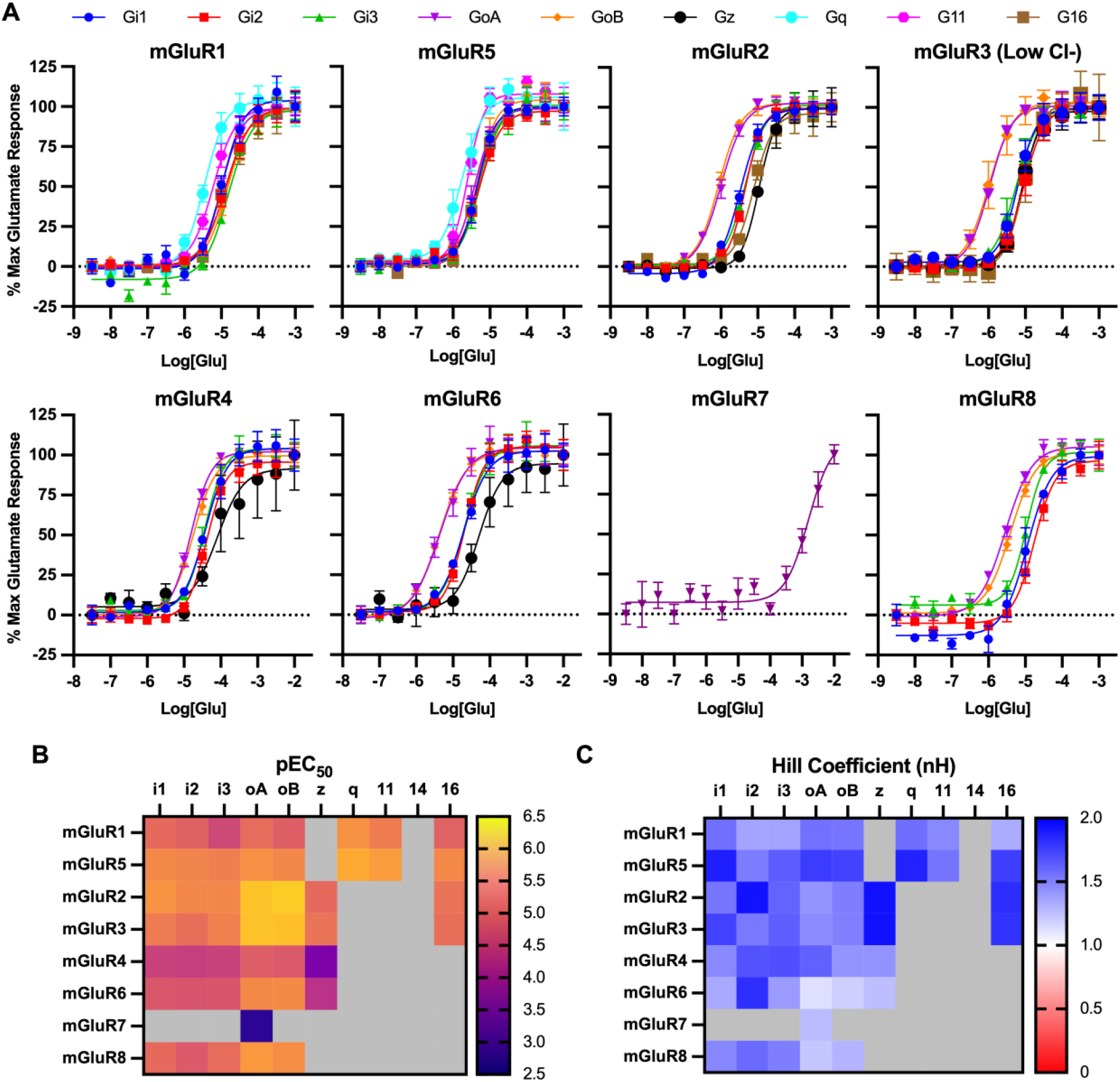
Glutamate dose response curves illustrating relative efficacy and potency of responses of each mGluR through each responding G protein. A, glutamate dose response curves for the indicated G⍺ proteins when coexpressed with mGluRs1-8. Note that mGluR7 only showed responses to G_oA_, and only at glutamate concentrations above 1 mM, so accurate efficacy and potency estimates were not possible. Heat maps are also shown, illustrating calculated EC_50_ values (B) and Hill coefficients (C) for the indicated mGluR homodimer with each responding G⍺ protein in the NanoBRET assay.

## Discussion

### G protein coupling profiles of mGluR homodimers

We show here for the first time a comprehensive mGluR - G protein coupling profiling assessment with every homodimeric member of the human mGluR family against representatives of each G protein family. Heat maps summarizing all of the maximal responses and activation kinetics are shown in Fig. 7A&B, respectively. These data show that the group I mGluRs, mGluR1 and mGluR5, couple to both the G_q/11_ and G_i/o_ proteins, as previously suggested (*2, 4*). No evidence for group I mGluR coupling to members of the G_s_ family was seen, including in the presence of the mGluR5 PAM VU042 (*8*). In addition, the group II and III mGluRs coupled almost exclusively to the G_i/o_ proteins, with the only exception being a weak, slow activation of the promiscuous G_16_ by mGluR2 and to a lesser degree mGluR3. While G protein profiling has been examined to some extent on representative members of the mGluR family, to our knowledge this is the first comprehensive assessment of all of the mGluRs with a large set of G⍺ proteins. One recent study examined G protein profiles of many GPCRs using an effector translocation based BRET assay and reported results on mGluR2, 4, 5, 6, and 8 largely consistent with those reported here (*18*). One notable exception was a reported coupling between mGluR5 and G⍺_14_, which we did not see. However, we would note that in that study, the authors similarly did not observe coupling of mGluR5 to members of the G⍺_s_ family. Our results were consistent with that finding.

**Figure 7.**
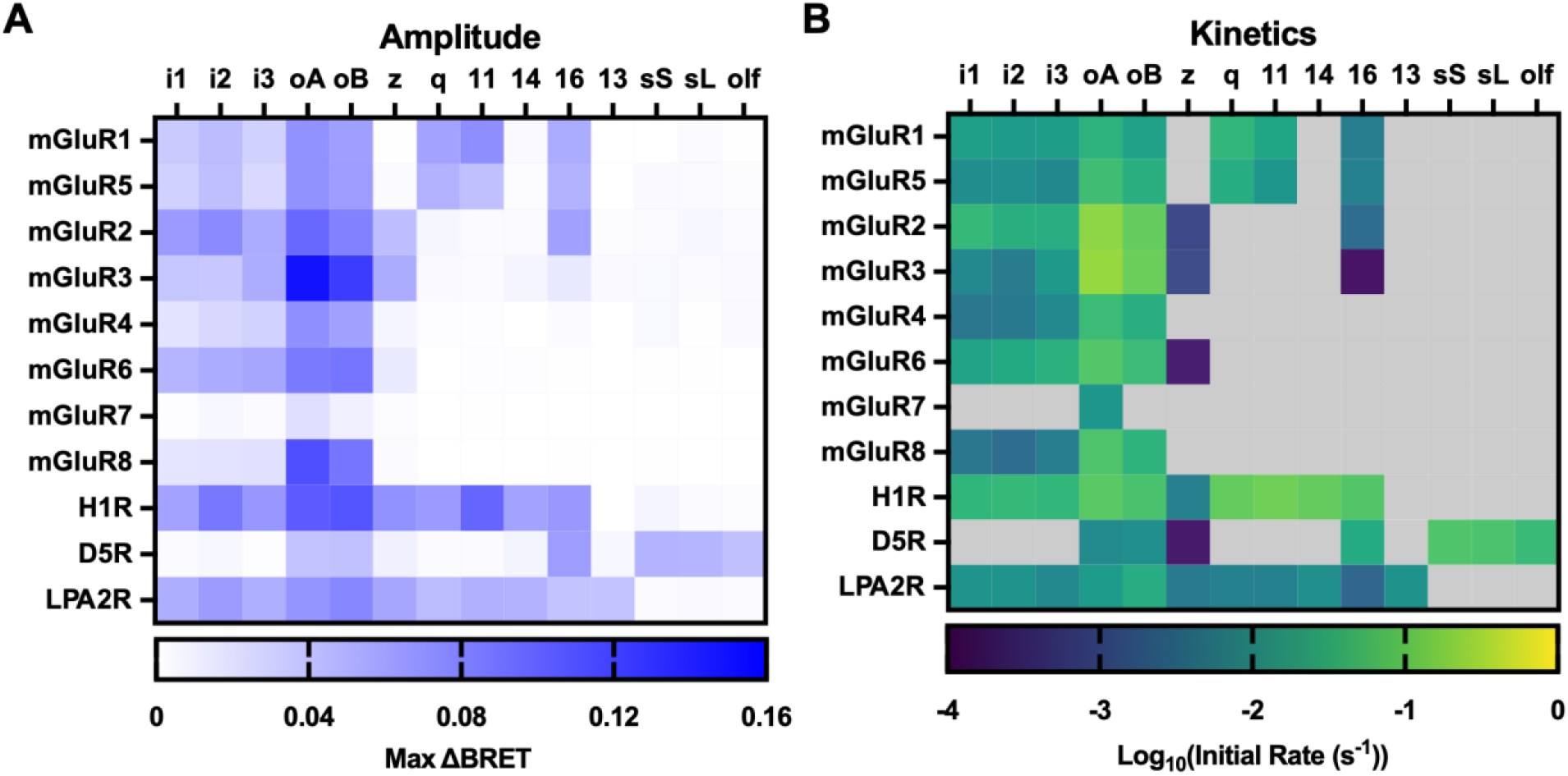
Summary of mGluR-G protein responses of all mGluR homodimers. A, Summary of signal amplitude data displayed as raw ΔBRET according to blue intensity scale shown below for each mGluR-G protein pair. B, Initial rate data displayed as Log_10_ transformed initial rates according to the color scale shown below for each mGluR-G protein pair. Heatmaps in both A&B display the average value of three biologic replicates. Data are from the same experimental replicates shown in Figs. 2-5.

### G protein activation kinetics

Comparing signaling of each of the members of the mGluR family, it is apparent that the efficacies in the NanoBRET assay are somewhat comparable. However, examination of the kinetics reveals that the initial rates of activation of different mGluR/G protein pairings to be quite variable (Fig. 7). Under some circumstances such as desensitization or differences in receptor expression level, maximal efficacy in this kind of assay may be misleading. Thus, activation kinetics can provide a more objective assessment of the efficiency of receptor-G protein coupling (*32*). In general, we see the Group II mGluRs are highly efficient receptors, activating G proteins at considerably faster rates compared to the Group I or Group III receptors (Fig 7). The only major exception to this trend is with G_16_ signaling, where the bona-fide G_q/11_ coupling mGluR1 and mGluR5 show faster activation than the Group II receptors. Additionally, most mGluRs showed faster kinetics through G_o_ proteins than other G⍺ proteins, with only mGluR1 showing slightly faster kinetics through G_q_. Although this finding is going to be largely influenced by the affinity of each individual G⍺ protein for the Gβ*γ* used, receptor level effects are clearly present given the H1R was able to activate G_q/11_ proteins with faster kinetics than the G_o_ protein, and the D5R was able to activate G_s_ proteins with faster kinetics than G_o_ proteins. These additional findings suggest that mGluRs favor signaling through G_o_ over other G⍺_i/o_ family members.

Regarding GPCR-G protein coupling experiments, especially when using activation kinetics as a proxy for coupling efficiency, it is important to consider the expression of the Regulators of G protein Signaling (RGS) proteins in the cells assayed. This is important because while RGS proteins facilitate deactivation kinetics of G⍺ proteins by acting as GTPase activating Proteins (GAPs), they can also accelerate activation kinetics (*33–36*). HEK293 cells have been suggested to express a wide array of RGS proteins (*37*), but more recent work using RNA microarrays suggested that the expression may be more limited (*38*). Still, while RGS protein expression may affect interpretation of specific details such as kinetics, it is unlikely that a different compliment of RGSs will yield coupling to a specific G⍺ protein in another system where none was observed here. It is also unlikely that the rank order of coupling efficiency of receptors would be altered with different RGSs due to them exerting their effects on the G protein level rather than the receptor level. For example, we observed coupling to G⍺_oA_ through the group II mGluRs to be more efficient than through the Group III receptors. Since they all couple to the same family of G proteins, it is unlikely this relation will be different with other RGSs that also act as GAPs through these same G proteins.

These data highlight an interesting aspect of mGluR-G protein coupling across the family, specifically coupling efficiency. We found that the group II mGluRs exhibited the fastest signaling kinetics when coupled to G_i/o_ proteins and mGluR2 in particular when coupled to all members of the G_i/o_ family (Fig. 7). These results suggest that in the physiological context, when mGluRs reside in the synaptic environment and are likely to be exposed to saturating concentrations of glutamate for only brief periods of time, that the group II mGluRs may play a dominant role in the modulation of synaptic transmission. Another interesting aspect of mGluR-G⍺ protein coupling is the differences in potency that the receptors activate different G⍺ proteins. These differences probably reflect a combination of varying affinities that each G⍺ protein has with the active state of the receptors and the abundance of each G⍺ in cell. The group II and III receptors show a clear preference for the G⍺_o_ proteins, followed by G⍺_i1-3_, with most also activating G⍺_z_. By contrast, the group I receptors activate the G⍺_q/11_ proteins with the highest potency, followed by members of the G_i/o_ family with relatively similar potencies.

In this study, we examine mGluR activation kinetics. One recently published study examined intradimer conformational changes of several mGluR dimers using a FRET assay (*39*). There, authors reported that glutamate induced changes in FRET were measurable for 5 of the 8 mGluR homodimers. Interestingly and seemingly at odds with the kinetics of G protein activation described here, they reported that the fastest on kinetics were associated with mGluR1, and the slowest with mGluR2. However, it should be noted that in that study, what was measured was the movement of the subunits within each dimer that would lead to activation, while we measured the presence of active, ‘free’ Gβ*γ*, which can be considered a measure of the efficiency of guanine nucleotide exchange of each active receptor, not the kinetics of the conformational changes of an inactive receptor transitioning to an active one. Comparing these values directly, inactive to active conformational changes of even the slowest receptor in that study was on the order of 10-20 msec (*39*), still several orders of magnitude faster than the rates of guanine nucleotide exchange of all of the receptors in this study with which coupling was detected. Thus from a physiological perspective, it is still reasonable to consider the group II mGluRs as the most efficient activators of G proteins in the family.

## Materials and Methods

### Plasmids and Molecular Biology

Plasmids encoding G_i1_, G_i2_, G_i3_, G_oA_, G_sS_, Ric8B, mGluR6, and mLPA2R were gifts from Dr. Cesare Orlandi (University of Rochester) (*40*). Plasmids encoding G_oB_, G_z_, G_q_, G_11_, G_14_, G_13_, G_sL_, G_olf_, Gβ1, G*γ*2, masGRK3ct, EGFP-PTX-S1 and the D5R were gifts from Dr. Stephen Ikeda. (NIAAA) The pmVenus-N1 plasmid was a gift Dr. Steven Vogel (NIAAA). The CMV-hEAAT3 plasmid was a gift from Susan Amara (NIMH; Addgene plasmid #32815). The pH1R-P2A-mCherry-N1 was a gift from Dorus Gadella (Addgene plasmid # 84330) (*41*). A plasmid encoding for NanoLuc was a gift from Dr. John Lueck (University of Rochester) (*42*). All plasmids were verified by full sanger sequencing before use.

The Gβ*γ*-masGRK3ct sensor components, mVenus(156-239)-Gβ1, mVenus(1-155)-G*γ*2, and masGRK3ct-NL were assembled to be identical to those previously reported (*20*), with the exception of replacing RLuc8 with NLuc. For masGRK3ct-NLuc, the previously assembled masGRK3ct constructs (amino acids 495-688 of bovine GRK3 with the myristic acid sequence, MGSSKSKTSNS added to the N-terminus) and NanoLuc were copied from their original plasmids with PCR with appropriate overhangs for Gibson assembly into the EcoRV site in pCDNA3.1(+). A GCCACC Kozak sequence was added before the start codon of masGRK3ct. A GGG linker was incorporated into both overhang and both fragments. Next, pCDNA3.1(+) was digested with EcoRV, and the digested pCDNA3.1(+) was added along with the masGRK3ct and NanoLuc PCR products into an NEBuilder reaction. The reaction product was then transformed into XL10-Gold Ultracompetent E.Coli cells and colonies were screened for successful assembly. The mVenus(156-239)-Gβ1 and mVenus(1-155)-G*γ*2 were cloned with an identical procedure with the incorporation of GGSGGG linker in the overhangs between the mVenus fragments and protein.

The human mGluR constructs (except mGluR6) were synthesized by GenScript in fragments and assembled in lab. Each hmGluR coding sequences was domesticated by eliminating all BsaI, BbsI, BsmBI, SapI, and AarI restriction sites by introducing silent mutations. Each coding sequence was then divided into two, with a GCCACC Kozak sequence being added to the first fragment immediately before the start codon, the native stop codon was changed to a TGA stop, and overhangs were added for Gibson assembly into the EcoRV site of pCDNA3.1(+). Each fragment was synthesized by GenScript, and once received, the appropriate fragments were mixed with EcoRV digested pCDNA3.1(+) and subjected to a NEBuilder reaction.

### HEK293T cell culture and transfection

HEK293T cell cultures were maintained in growth media consisting of Dulbecco’s Modified Eagle Media (DMEM) supplemented with 10% fetal bovine serum (FBS), 2 mM L-alanyl-L-glutamine (1x GlutaMax), 100 units/mL penicillin, and 100 μg/mL streptomycin at 37°C in 5% carbon dioxide. Cells were routinely harvested, counted, and replated every 2-3 days to prevent cultures from overgrowing. Prior to counting, cells were resuspended in Trypan Blue stain to allow for assessment of cell death, which was routinely under 10%. When conducting experiments cells were plated as described in the individual assay protocols in growth media 4 hours prior to transfection. To transfect cells, cDNA was combined polyethylenimine (PEI) in unsupplemented DMEM for 20 minutes before addition to cells. The amount of PEI added was adjusted based on the total amount of cDNA used for the transfection, using 4μL of 7.5 mM PEI per 1μg of cDNA. For some assays, the media on cells was changed to DMEM supplemented with 2% FBS only immediately prior to addition of the transfection mixture.

### NanoBRET experiments using the Gβ*γ*-masGRK3ct sensor

For NanoBRET experiments, a modified standard protocol based on previously published protocols (*21*), or the mGluR optimized protocol described here were conducted. The day before the assay, HEK293T cells were plated into 6 well plates at 2 million cells per well in 1.5 mL of growth media. Four hours after plating transfections containing 200 ng masGRK3ct-NL, 200 ng mVenus(156-239)-Gβ_1_, 200 ng mVenus(1-155)-G*γ*_2_, 400 ng EAAT3, 600 ng G⍺ protein, and 800 ng of receptor was assembled in 500 uL of supplemented DMEM with an appropriate amount of PEI. After 20 minutes, the transfection mixture was added to the cells dropwise. If the mGluR optimized protocol was being used, the media on the cells was changed to 1.5 mL of DMEM with 2% FBS immediately prior to adding the transfection. Cells were then allowed to transfect overnight.

The following day, the cell media was removed, and the wells were washed once with phosphate buffered saline (PBS) with no calcium or magnesium. The PBS was then aspirated and PBS with 5 mM EDTA was added. Cells were then incubated in PBS with EDTA at 37°C for 5 minutes. Cells were then harvested by titration and collected into microcentrifuge tubes. Cells were then pelleted and washed three times with imaging buffer consisting of 136 mM NaCl, 560 μM MgCl_2_, 4.7 mM KCl, 1 mM Na_2_HPO4, 1.2 mM CaCl_2_, 10 mM HEPES, and 5.5 mM Glucose.

Experiments conducted with Low Cl-buffer used the same imaging buffer except the NaCl was replaced with 136 mM sodium gluconate. After the third wash, the cells were resuspended in appropriate volume of imaging buffer and 25 μL of cells were transferred into each well of opaque, flat bottom, white 96-well plate. For the standard procedure, cells were than assayed immediately. For the mGluR optimized protocol, cells were allowed to incubate in the plate for 1 hour before being assayed.

Cell responses were assayed using a PolarStar Omega multimodal plate reader (BMG Labtech) equipped with dual emission PMTs and two compound injectors. To select for NanoLuc and mVenus light, a 485/15 and a 535/30 filters were used respectively. Luminescent signals were integrated for 200 ms time bins, with the gain for both detectors set to 2000. Injectors were loaded with either NanoGlo reagent (Promega, 1:250 dilution in imaging buffer) or test compounds and addition was automated by the plate reader. Injections were done at a speed of 430 μL per sec. For kinetic dose response experiments, 60 second time courses were used with 25 μL of NanoGlo being injected at -19 seconds and 50 μL of 2x test compound being added at t = 0 seconds. For G⍺ profiling experiments, 120 second time courses were used, with injections at the same time points. Experiments reported in Figs. 1-5 were conducted in kinetic mode, as illustrated. Dose-response experiments reported in Fig. 6 were mainly conducted in endpoint mode (NanoGlo reagent, then glutamate added manually and data points recorded subsequently for ∼90 sec. for most G proteins, or ∼210 sec for G⍺_z_ and G⍺_16_ to account for slower activation kinetics). The exception was the experiments involving the group I mGluRs which were acquired in kinetic mode due to the acute desensitization that was particularly evident with mGluR5 (see Fig. 2).

### Data Analysis

For BRET experiments, the BRET ratio was calculated by dividing the mVenus signals (luminescence in the 535 channel) by the NanoLuc signals (luminescence in the 485 channel):

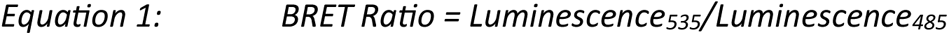

Reponses are analyzed as ΔBRET which is the average basal BRET ratio subtracted from the average stimulated BRET ratio:

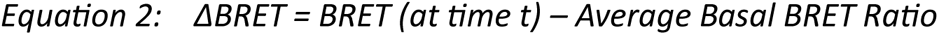

For endpoint assays, all 3 basal reading were averaged for the average basal BRET ratio, and all 5 readings post stimulation were average for the average stimulated BRET ratio. For kinetic experiments the basal BRET ratio was calculated as the average BRET ratio for 5 seconds immediately prior to test compound injection. For the average stimulated BRET ratio, the average BRET ratio for the last 10 seconds of the trace was used for non-desensitizing signals. For desensitizing signals, the average BRET ratio of a 10 second window centered at the signal’s peak was used.

To analyze the kinetics of BRET curves, the upstroke of each response was fit in GraphPad Prism (v. 9.3.1) using 1 of 2 models. The first model used was Pharmechanics’s “Baseline then rise to steady state time course” equation (*32*) (a single-phase exponential association model):

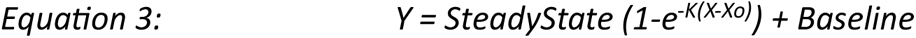

where Y is the response, SteadyState is plateau of the response, K is the rate constant, Baseline is the average baseline of the signal, and X_0_ is the time the response initiates, and X is time. The second model used was a custom programmed two-phase exponential association model based on the above equation:

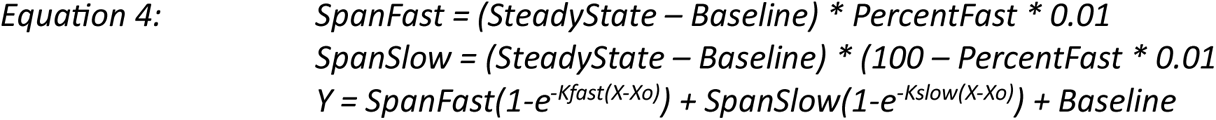

where Y is the response, SteadyState is plateau of the response, K_fast_ is the faster rate constant, K_slow_ is the slower rate constant, Baseline is the average baseline of the response, PercentFast is the percent contribution of the fast component to the response, X_0_ is the time the response initiates, and X is time. Calculation of the initial rates was then conducted by multiplying the SteadyState by the rate constant K for single association curves:

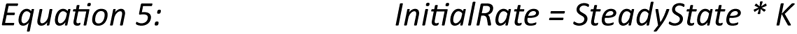

or by multiplying the SteadyState by the weighted average of K_fast_ and K_slow_ for two-phase associations:

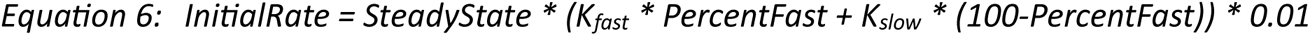

To analyze dose responses, data was imported into GraphPad Prism. Individual dose responses were fitted with the built in four parameter logistic equation:

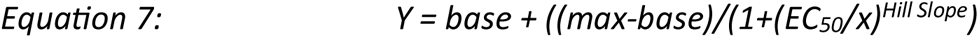

Where Y is the response, base is the response baseline, max is the maximum response (E_max_), EC_50_ is the concentration of drug that produces a 50% response. To aggregate responses from multiple replicates from the same conditions, the fit parameters were copied to Microsoft Excel and average values and standard errors were calculated for each condition.

### Statistics

All statistical analysis was conducted in GraphPad Prism. Results are reported in the figure legends in (P = [P-value]) format. Details of the test (type of test, results, significance levels) are indicated in the figure legend.

## Acknowledgments

We thank Dr. Cesare Orlandi (University of Rochester) for use of his plate reader, and for helpful guidance. These studies were supported by grants R21NS126779, R03NS124987, and R01MH125849 to PJK.

## References

1. K. J. Gregory, Asymmetry is central to excitatory glutamate receptor activation. Nat Struct Mol Biol 28, 633–635 (2021).

2. P. J. Kammermeier, S. R. Ikeda, Expression of RGS2 alters the coupling of metabotropic glutamate receptor 1a to M-type K+ and N-type Ca2+ channels. Neuron 22, 819–829 (1999).

3. C. Avet et al., The PAR2 inhibitor I-287 selectively targets Galpha(q) and Galpha(12/13) signaling and has anti-inflammatory effects. Commun Biol 3, 719 (2020).

4. B. A. McCool et al., Rat group I metabotropic glutamate receptors inhibit neuronal Ca2+ channels via multiple signal transduction pathways in HEK 293 cells. J Neurophysiol 79, 379–391 (1998).

5. A. S. Hauser et al., Common coupling map advances GPCR-G protein selectivity. Elife 11, (2022).

6. R. K. Senter et al., The Role of mGlu Receptors in Hippocampal Plasticity Deficits in Neurological and Psychiatric Disorders: Implications for Allosteric Modulators as Novel Therapeutic Strategies. Curr Neuropharmacol 14, 455–473 (2016).

7. Y. Kubo, M. Tateyama, Towards a view of functioning dimeric metabotropic receptors. Curr Opin Neurobiol 15, 289–295 (2005).

8. C. Nasrallah et al., Direct coupling of detergent purified human mGlu(5) receptor to the heterotrimeric G proteins Gq and Gs. Sci Rep 8, 4407 (2018).

9. I. Sebastianutto et al., D1-mGlu5 heteromers mediate noncanonical dopamine signaling in Parkinson’s disease. J Clin Invest 130, 1168–1184 (2020).

10. M. Tateyama, Y. Kubo, Coupling profile of the metabotropic glutamate receptor 1alpha is regulated by the C-terminal domain. Mol Cell Neurosci 34, 445–452 (2007).

11. C. M. Niswender, P. J. Conn, Metabotropic glutamate receptors: physiology, pharmacology, and disease. Annu Rev Pharmacol Toxicol 50, 295–322 (2010).

12. C. Upreti, X. L. Zhang, S. Alford, P. K. Stanton, Role of presynaptic metabotropic glutamate receptors in the induction of long-term synaptic plasticity of vesicular release. Neuropharmacology 66, 31–39 (2013).

13. L. Mao, M. Guo, D. Jin, B. Xue, J. Q. Wang, Group III metabotropic glutamate receptors and drug addiction. Front Med 7, 445–451 (2013).

14. Y. Nakajima et al., Molecular characterization of a novel retinal metabotropic glutamate receptor mGluR6 with a high agonist selectivity for L-2-amino-4-phosphonobutyrate. J Biol Chem 268, 11868–11873 (1993).

15. P. J. Conn, J. P. Pin, Pharmacology and functions of metabotropic glutamate receptors. Annu Rev Pharmacol Toxicol 37, 205–237 (1997).

16. N. Okamoto et al., Molecular characterization of a new metabotropic glutamate receptor mGluR7 coupled to inhibitory cyclic AMP signal transduction. J Biol Chem 269, 1231–1236 (1994).

17. C. H. Habrian et al., Conformational pathway provides unique sensitivity to a synaptic mGluR. Nature communications 10, 5572 (2019).

18. C. Avet et al., Effector membrane translocation biosensors reveal G protein and betaarrestin coupling profiles of 100 therapeutically relevant GPCRs. Elife 11, (2022).

19. A. Ritzen, J. M. Mathiesen, C. Thomsen, Molecular pharmacology and therapeutic prospects of metabotropic glutamate receptor allosteric modulators. Basic Clin Pharmacol Toxicol 97, 202–213 (2005).

20. B. Hollins, S. Kuravi, G. J. Digby, N. A. Lambert, The c-terminus of GRK3 indicates rapid dissociation of G protein heterotrimers. Cell Signal 21, 1015–1021 (2009).

21. I. Masuho, K. A. Martemyanov, N. A. Lambert, Monitoring G Protein Activation in Cells with BRET. *Methods in molecular biology (Clifton*, N.J.) 1335, 107–113 (2015).

22. V. Hlavackova et al., Sequential inter- and intrasubunit rearrangements during activation of dimeric metabotropic glutamate receptor 1. Sci Signal 5, ra59 (2012).

23. C. Goudet et al., Heptahelical domain of metabotropic glutamate receptor 5 behaves like rhodopsin-like receptors. Proc Natl Acad Sci U S A 101, 378–383 (2004).

24. V. Hlavackova et al., Evidence for a single heptahelical domain being turned on upon activation of a dimeric GPCR. EMBO J 24, 499–509 (2005).

25. P. J. Kammermeier, S. R. Ikeda, Desensitization of group I metabotropic glutamate receptors in rat sympathetic neurons. J Neurophysiol 87, 1669–1676 (2002).

26. G. K. Dhami, S. S. Ferguson, Regulation of metabotropic glutamate receptor signaling, desensitization and endocytosis. Pharmacol Ther 111, 260–271 (2006).

27. S. R. Ikeda, S. W. Jeong, P. J. Kammermeier, V. Ruiz-Velasco, M. M. King, Heterologous expression of a green fluorescent protein-pertussis toxin S1 subunit fusion construct disrupts calcium channel modulation in rat superior cervical ganglion neurons. Neurosci Lett 271, 163–166 (1999).

28. A. S. Tora et al., Chloride ions stabilize the glutamate-induced active state of the metabotropic glutamate receptor 3. Neuropharmacology 140, 275–286 (2018).

29. P. J. Kammermeier, Constitutive activity of metabotropic glutamate receptor 7. BMC Neurosci 16, 17 (2015).

30. Y. Wang et al., The GABA(B) receptor mediates neuroprotection by coupling to G(13). Sci Signal 14, eaaz4112 (2021).

31. N. Abreu, A. Acosta-Ruiz, G. Xiang, J. Levitz, Mechanisms of differential desensitization of metabotropic glutamate receptors. Cell reports 35, 109050 (2021).

32. S. R. J. Hoare, P. H. Tewson, A. M. Quinn, T. E. Hughes, L. J. Bridge, Analyzing kinetic signaling data for G-protein-coupled receptors. Sci Rep 10, 12263 (2020).

33. C. A. Doupnik, N. Davidson, H. A. Lester, P. Kofuji, RGS proteins reconstitute the rapid gating kinetics of gbetagamma-activated inwardly rectifying K+ channels. Proc Natl Acad Sci U S A 94, 10461–10466 (1997).

34. O. Saitoh, Y. Kubo, Y. Miyatani, T. Asano, H. Nakata, RGS8 accelerates G-protein-mediated modulation of K+ currents. Nature 390, 525–529 (1997).

35. S. W. Jeong, S. R. Ikeda, Differential regulation of G protein-gated inwardly rectifying K(+) channel kinetics by distinct domains of RGS8. J Physiol 535, 335–347 (2001).

36. J. J. Choi et al., Protein trans-splicing and characterization of a split family B-type DNA polymerase from the hyperthermophilic archaeal parasite Nanoarchaeum equitans. J Mol Biol 356, 1093–1106 (2006).

37. G. Laroche, P. M. Giguere, B. L. Roth, J. Trejo, D. P. Siderovski, RNA interference screen for RGS protein specificity at muscarinic and protease-activated receptors reveals bidirectional modulation of signaling. Am J Physiol Cell Physiol 299, C654–664 (2010).

38. B. K. Atwood, J. Lopez, J. Wager-Miller, K. Mackie, A. Straiker, Expression of G protein-coupled receptors and related proteins in HEK293, AtT20, BV2, and N18 cell lines as revealed by microarray analysis. BMC Genomics 12, 14 (2011).

39. T. Kukaj, C. Sattler, T. Zimmer, R. Schmauder, K. Benndorf, Kinetic fingerprinting of metabotropic glutamate receptors. Commun Biol 6, 104 (2023).

40. I. Masuho et al., Distinct profiles of functional discrimination among G proteins determine the actions of G protein-coupled receptors. Sci Signal 8, ra123 (2015).

41. J. van Unen et al., Quantitative Single-Cell Analysis of Signaling Pathways Activated Immediately Downstream of Histamine Receptor Subtypes. Mol Pharmacol 90, 162–176 (2016).

42. W. Ko, J. J. Porter, M. T. Sipple, K. M. Edwards, J. D. Lueck, Efficient suppression of endogenous CFTR nonsense mutations using anticodon-engineered transfer RNAs. Mol Ther Nucleic Acids 28, 685–701 (2022).

