## Supplemental Information for "Signaling specificity and kinetics of the human metabotropic glutamate receptors"

Figure S1. Signaling profiles of control receptors. Data are displayed as in Figures 2-5. A-C, illustrates responses of the H1 receptor (100 μM histamine) to each Gα protein tested, including a robust response to $G_{14}$. D-F, shows responses to the LPA2R (10 μM lysophosphatidic acid), which responded to $G_{13}$ as well as G proteins in the $G_{i/o}$ and $G_{q/11}$ families. G-I, illustrates D5 receptor responses (100 μM dopamine), which showed responses to each member of the $G_s$ family tested. Stimuli (respective agonists) were delivered at time = 0. The dark solid line indicates the average response of 3 biologic replicates and the shaping indicates the SEM. Initial rates are displayed as the average ± SEM.

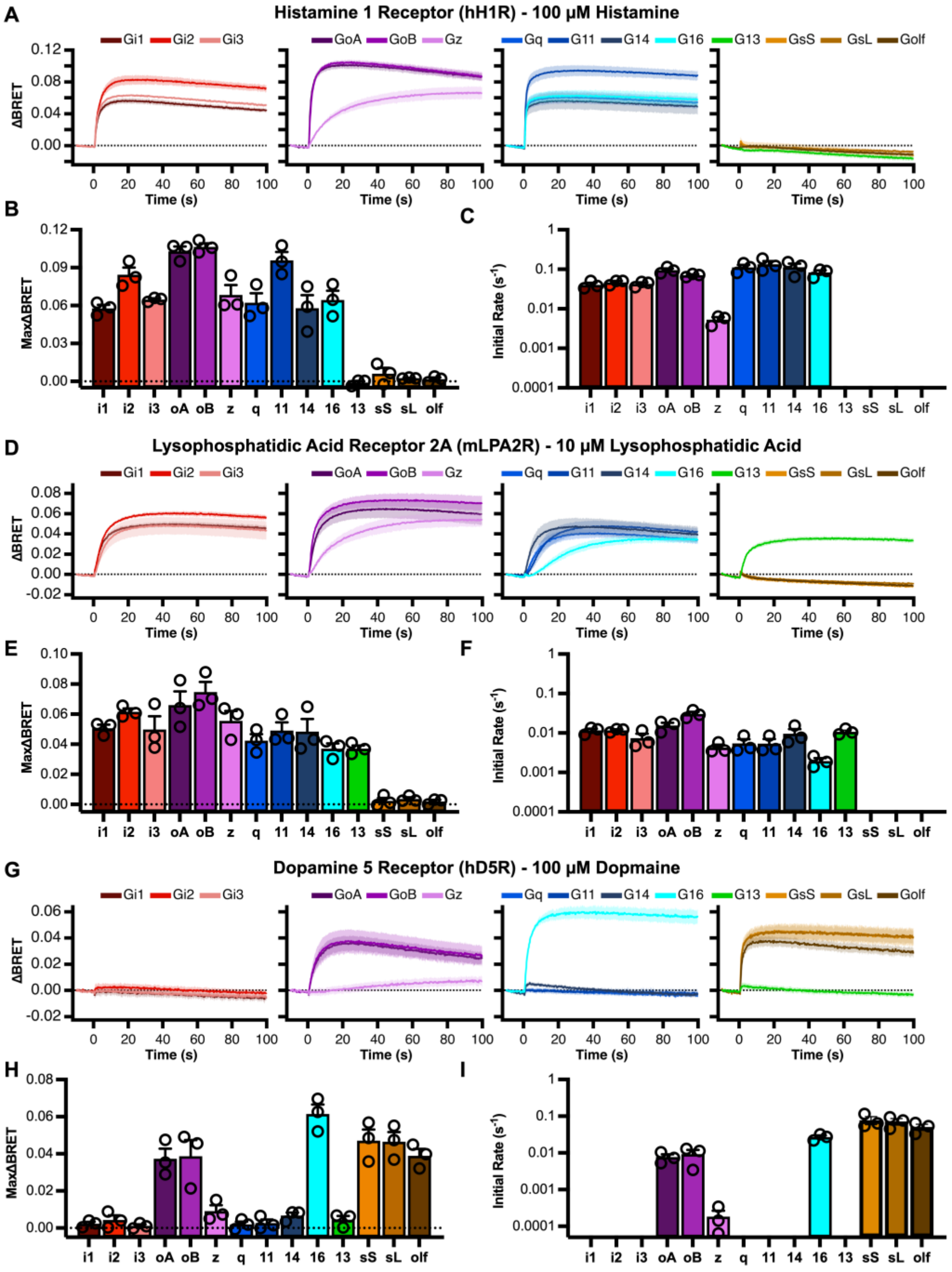

Figure S2. The mGluR5 PAM VU0424 does not induce $G_s$ coupling. Average responses of mGluR5 in the NanoBRET system to 1 mM glutamate alone (A), 10 µM VU0424 (B), and both glutamate and VU0424 (C). D shows the maximum deltaBRET responses to each condition ±SEM. E shows the average initial rates. Individual responses to glutamate (○), VU0424 (□), and both drugs (◊) are shown as the indicated symbols.

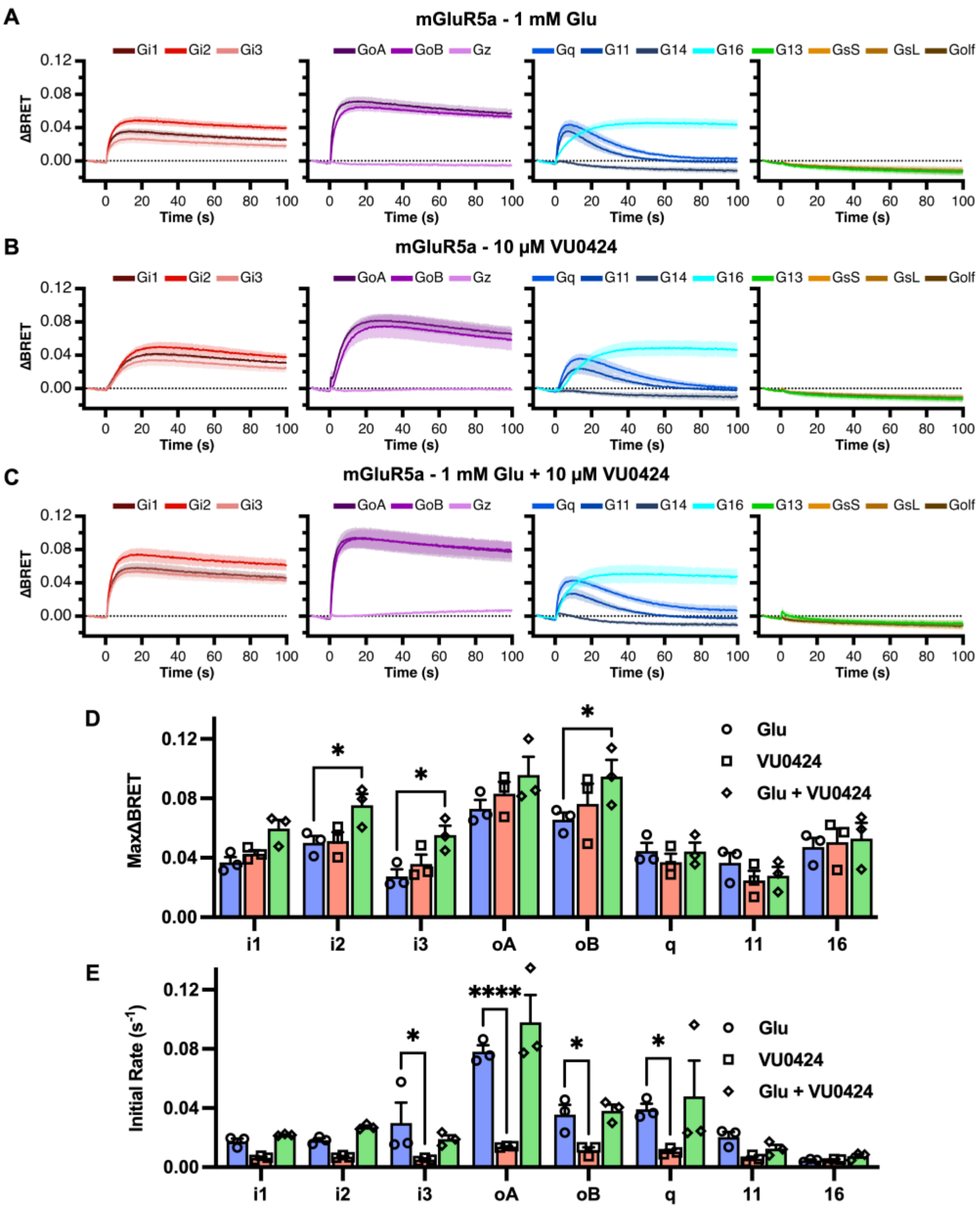

Figure S3. Low $Cl^-$ does not alter mGluR2 signaling profiles. A&B illustrate kinetic responses of mGluR2 activation of 14 Gα proteins in normal (A) and low $Cl^-$ (B) external solutions. Summary data of the amplitudes and initial rates of hmGluR2 signals in High Cl- buffer (gray) and Low Cl- buffer (purple) are also shown in C and D, respectively. Bars indicate the average ± SEM. Statistics indicate the results of a two-way ANOVA with Holm-Šídák *post hoc* test. * = P<0.05, ** = P<0.005, *** = P<0.0005, and **** = P<0.0001 between the indicated conditions. Individual comparison of amplitudes did not reach statistical significance for any condition. E, Correlation of signal amplitudes in normal (high) $Cl^-$ buffer and Low $Cl^-$ buffer. F, Correlation of signal initial rates in High $Cl^-$ and Low $Cl^-$ buffer. Data points show average ± SEM. Linear regression lines are displayed as a solid black line, with the 95% confidence interval being indicated by the dashed line. A theoretical line with slope 1 and intercept of (0,0) is shown as a dashed gray line for reference.

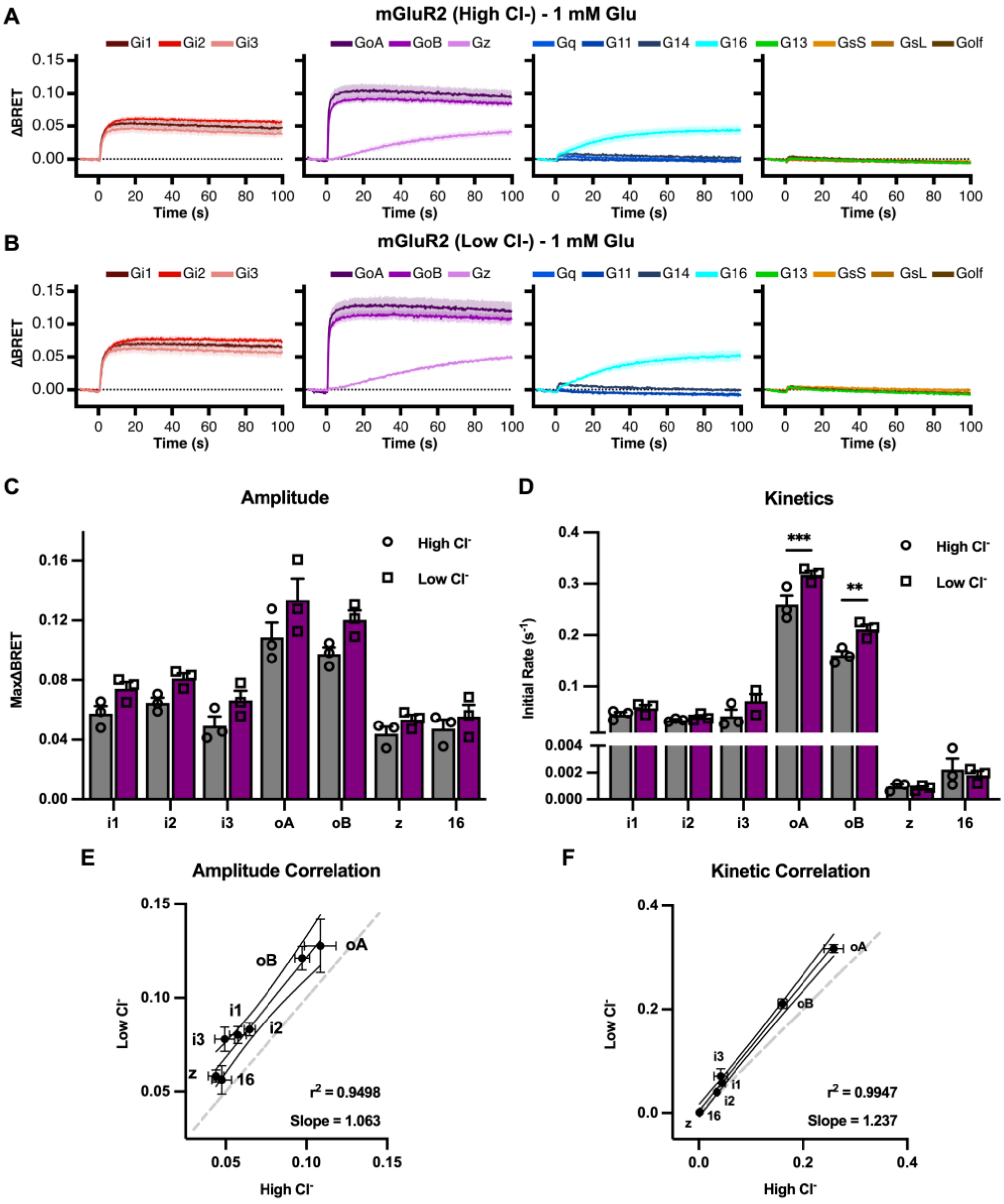

Figure S4. Low $Cl^-$ solution right shifts glutamate efficacy for every mGluR homodimer tested. A, Glutamate dose responses collected from the same mGluR1a-Gq, mGluR2-GoA, mGluR4-GoA, mGluR5a-Gq, and mGluR8-GoA expressing cells in either High $Cl^-$ buffer (black) or low $Cl^-$ buffer (purple). N = 3-5 biologic replicates per condition. Individual responses are displayed as the average ± SEM. B and C show summaries of the potency and efficacy changes that accompanied changes in extracellular $Cl^-$ in response to 1 mM glutamate for mGluRs 1, 2, 4, 5, and 8 with the indicated Gα protein co-expressed. Panel D shows glutamate dose responses for mGluR2 with each Gα protein in low $Cl^-$ solution (compare to Fig. 5). E shows a correlation between the glutamate $pEC_{50}$ values measured for mGluR2 with each Gα protein in low (y-axis) and high (x-axis) $Cl^-$.

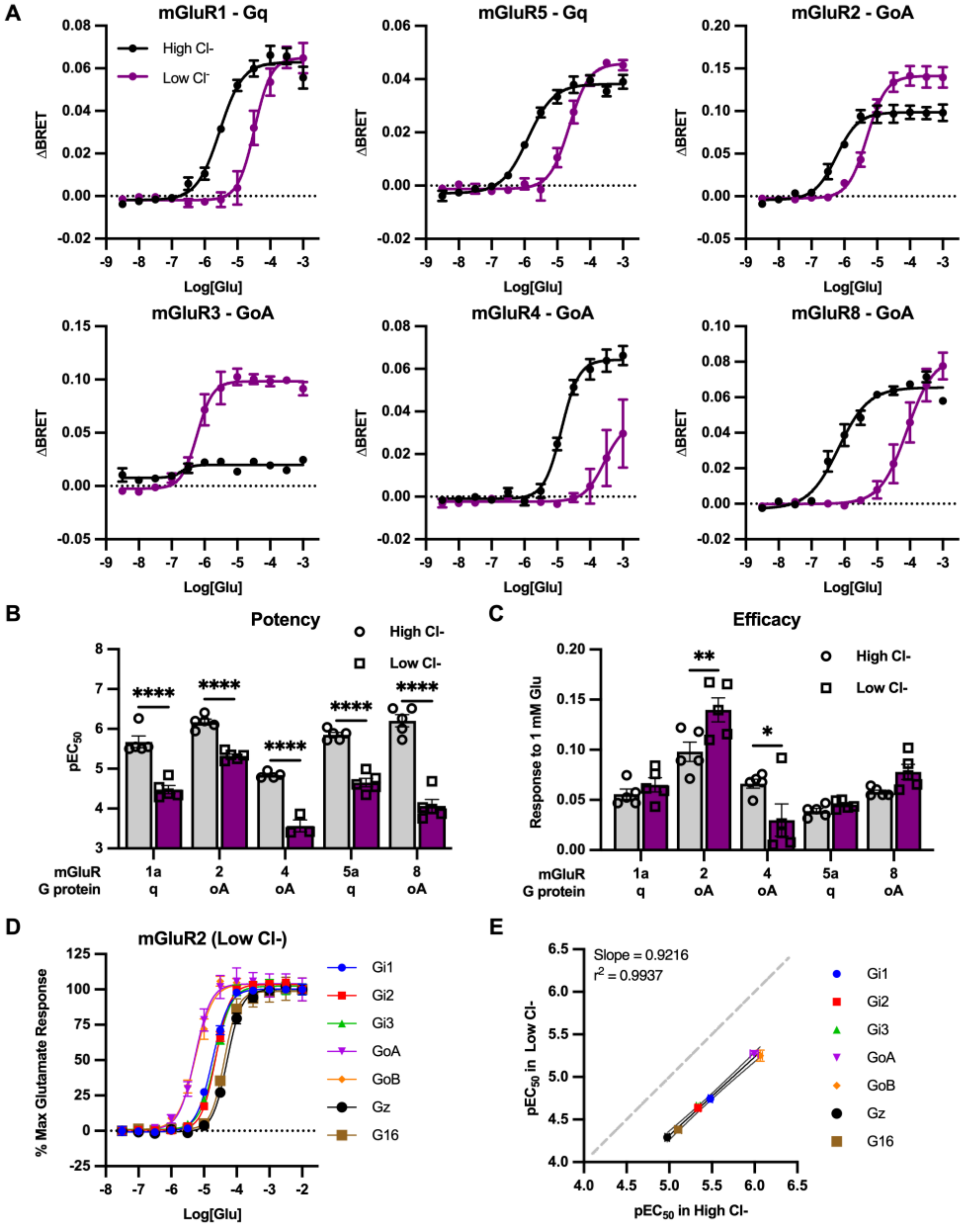
